# BacAnt: A Combination Annotation Server for Bacterial DNA Sequences to Identify Antibiotic Resistance Genes, Integrons, and Transposable Elements

**DOI:** 10.1101/2020.09.05.284273

**Authors:** Xiaoting Hua, Qian Liang, Min Deng, Jintao He, Meixia Wang, Wenjie Hong, Jun Wu, Bian Lu, Sebastian Leptihn, Yunsong Yu, Huan Chen

**Affiliations:** Department of Infectious Diseases, Sir Run Run Shaw Hospital, College of Medicine, Zhejiang University, Hangzhou, China; Key laboratory of microbial technology and bioinformatics of Zhejiang Province, Zhejiang Institute of Microbiology, Hangzhou 310012, China; Regional Medical Center for National Institute of Respiratory Diseases, Sir Run Run Shaw Hospital, School of Medicine, Zhejiang University, Hangzhou, China; NMPA Key laboratory for Testing and Risk Warning of Pharmaceutical Microbiology, Zhejiang Institute of Microbiology, Hangzhou 310012, China; Department of Infectious Diseases, The First Hospital of Jiaxing, The First Affiliated Hospital of Jiaxing University, Jiaxing 314000, China; Lin’an Center for Disease Control and Prevention, Lin’an, 311300, China; Xiaoshan Center for Disease Control and Prevention, Hangzhou, 311201, China; Zhejiang University-University of Edinburgh Institute, School of Medicine, Zhejiang University, Hangzhou, China

**Keywords:** Antimicrobial Resistance, Resistance genes, Gene Annotation, Insertion Sequences, IS, integron, Transposons, Tn

## Abstract

Whole genome sequencing (WGS) of bacteria has become a routine method in diagnostic laboratories. One of the clinically most useful advantages of WGS is the ability to predict antimicrobial resistance genes (ARGs) and mobile genetic elements (MGEs) in bacterial sequences. This allows comprehensive investigations of such genetic features but can also be used for epidemiological studies. A plethora of software programs have been developed for the detailed annotation of bacterial DNA sequences, such as RAST (Rapid Annotation using Subsystem Technology), Resfinder, ISfinder, INTEGRALL and The Transposon Registry. Unfortunately, to this day, a reliable annotation tool of the combination of ARGs and MGEs is not available, and the generation of genbank files requires much manual input. Here, we present a new webserver which allows the annotation of ARGs, integrons, and transposable elements at the same time. The pipeline generates genbank files automatically, which are compatible with easyfig for comparative genomic analysis. Our BacAnt code and standalone software package are available at https://github.com/xthua/bacant with an accompanying web application at http://bacant.net.

**Graphical Abstract:** 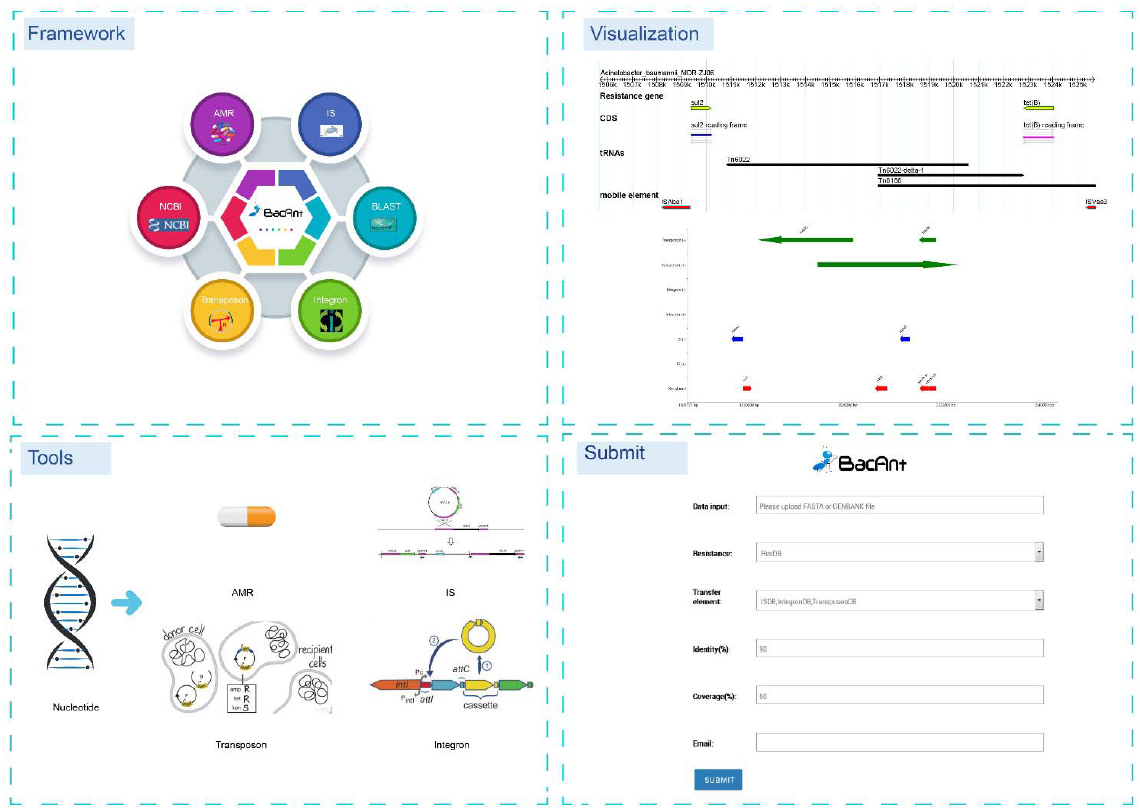

## Introduction

The era of next-generation sequencing (NGS) took off in 2005 with the commercial release of massively parallel pyrosequencing (Margulies et al., 2005). The NGS technology developed rapidly in the past years and has made substantial improvements in terms of quality and yield. With the rapid decrease of sequencing costs, falling by as much as 80% year over year, whole genome sequencing of bacteria has become a routine method in diagnostic laboratories (Didelot et al., 2012). NGS applications include whole genome sequencing (WGS), targeted NGS and metagenomic NGS. Among them, the most common use of WGS is for simultaneous identification, typing, and/or antimicrobial susceptibility prediction of pathogens (Mitchell and Simner, 2019). One of the most exciting advantages of NGS is the ability to predict antimicrobial resistance genes (ARGs) and mobile genetic elements (MGEs) in bacteria, which allows the investigation of both, the organization and structure of such genetic features, and the epidemiology for the distribution of bacterial strains or virulence genes, including the spread and distribution of antibiotic-resistant bacteria as part of surveillance programs (Zhou et al., 2015; Mitchell and Simner, 2019). Every day, a massive number of bacterial genomes is being sequenced using NGS technology in laboratories across the globe, with genomes released at remarkable rates. With this huge amount of data available, it is important to extract project-relevant information easily. However, in publicly available databases, most of bacterial genomes are available as contigs which have been constructed employing auto-annotation algorithms. Over the years, highly efficient methods for bacterial genome annotation have been developed that do not require much user input.

Rapid Annotation using Subsystem Technology (RAST) is a widely used webserver for genome annotations of microbial species (Aziz et al., 2008). Although the performance using RAST-based annotation is very useful, several important limitations remain. For example, RAST will label many Open Reading Frames (ORFs) as “hypothetical proteins”, and the performance to identify ARGs and label them as such, is fairly limited as the algorithm is not tailored towards this purpose. Based on the RAST system, the Pathosystems Resource Integration Center (PATRIC) improved the data collection of ARGs, and provided users a more powerful analysis for both genomes and individual genes (Wattam et al., 2018). Another available annotation server is Resfinder which is managed by the Center for Genomic Epidemiology; it provides a convenient way of identifying acquired ARGs in sequenced bacterial isolates (Zankari et al., 2012). In addition to annotations of ARGs, some databases specifically designed to annotate MGEs such as insertion sequences (ISfinder) and integrons (INTEGRALL) and transposable elements (The Transposon Registry) have been created (Siguier et al., 2006; Moura et al., 2009; Tansirichaiya et al., 2019). ISs are abundant mobile elements in bacteria, which are responsible for the mobilization of many genes, including those mediating ARG (Razavi et al., 2020). Such ARGs are often found in the genetic context of specific ISs, while ISs flanking regions are diverse (Razavi et al., 2020). For example, a clear association of ARGs with class 1 integrons can be observed (Partridge et al., 2018). The analysis of which ISs are associated with ARG genes would help to discover novel AMGs (Razavi et al., 2020). In addition, there is a major interest to explore how ARGs spread via MGEs (Che et al., 2021). The early identification of ARGs in bacteria would facilitate surveillance and molecular diagnostics (Razavi et al., 2020). Also, inter/intra-species genetic transfer events of MGEs are responsible for the emergence and rapid spread of resistance (Subedi et al., 2018). Therefore, the knowledge of MGE-associated drug resistance is crucial for the monitoring of resistance of microbial species. Unfortunately, up to now, a rapid annotation tool of the combination of ARGs and MGEs is not available, and the generation of genbank files has to be done manually. Therefore, we created a new program/pipeline called BacAnt, which rapidly and efficiently annotates ARGs, integrons, and transposable elements in a single step and generates a genbank file automatically which is compatible with the easyfig program for comparative genomic analysis.

## Materials and Methods

### Reference sequences

Three curated databases for BacAnt tool are used, including ResDB (resistance gene sequence database), IntegronDB (integron sequence database), and TransposonDB (transposon sequence database). We collected 5029 sequences from NCBI Bacterial Antimicrobial Resistance Reference Gene Database (PRJNA313047, https://www.ncbi.nlm.nih.gov/bioproject/PRJNA313047, version: 2019-09-06.1) into ResDB at 2019-12-01. In addition, we collected 1094 sequences from INTEGRALL (http://integrall.bio.ua.pt, version: 2017-11-30) to be included into IntegronDB by 2019-12-01. We also collected 234 sequences from THE TRANSPOSON REGISTRY (https://transposon.lstmed.ac.uk, version: 2019-07-23) into the TransposonDB by 2019-12-01.

### Program for identifying resistance genes and mobile elements

We created a python program (BacAnt) to identify resistance genes and mobile elements for bacteria nucleotide sequences with BLAST analysis. We first used the BLASTN program comparing input sequences with the reference database with an -evalue 10^−5^. For the detection of integrons, we used integron_finder to predict possible integrons and used the BLASTN program comparing the integron sequence with integronDB database for the best match sequence (Didelot et al., 2012). We then filtered the raw results by identities and coverage (blast align match length/subject length). All results that pass the identity and coverage filter are retained for further analysis. The default threshold was set to 90% for identities and to 60% for coverage. Finally, we display the filtered results in text and genbank format, while also providing a visual output. Three types of annotations for the sequences are displayed in the same figure to guide the analysis of the genomic sequence.

BacAnt has six parameters. The user has two choices regarding the input sequence file: --nucleotide (-n), fasta format or --genbank (-g), genbank format. The required output path: --resultdir (-o). --databases (-d), reference databases, select all by default. --coverages (-c), coverage threshold, 60% by default. --identities (-i), identities threshold, 90% by default. In average, it takes about 2 minutes for each run (number of available cores: 6; used threads: 24; memory 64G).

### Website for BacAnt

For the analysis to be performed online, we developed a website which we call http://bacant.net. The pipeline running on the server is based on Python/Django, which allows the user to upload sequence files for the rapid identification of ARGs and MGEs. The output format allows the display of graphic representations of the results. A demo report can be seen here: http://bacant.net/BacAnt/demo.

### Datasets for validation of BacAnt

BacAnt was validated with 1100 genomes (Table S1) from eight species (*A. baumannii, Bacillus cereus, Clostridioides difficile, E coli, Listeria monocytogenes, S. enterica, S. aureus, Vibrio parahaemolyticus*). The BacAnt output was analyzed and compared with the results of NCBI AMRFinder. The parameters of BacAnt used in the study was: identity 0.9, coverage 0.6.

### Examples analysis using BacAnt

Four genome sequences from NCBI, including *Acinetobacter baumannii* (2019.11.17), *Escherichia coli* (2019.5.7), *Salmonella enterica* (2019.6.4) and *Staphylococcus aureus* (2019.6.4) were used to illustrate the capabilities of our program. The accession number of the genomes used in this study were listed in Table S2 and Table S3. The number of drug-resistance genes and insertion sequences of each sample were identified in the BacAnt analysis and plotted as horizontal and vertical coordinates respectively. Scatter plots were created using the ggplot2 package in R (V3.6.2), together with the trend line (Wickham, 2016).

We created a network diagram by Cytoscape (v3.7.2) using the analysis results of BacAnt from four species genome sequences from NCBI (Shannon et al., 2003). Only resistance gene pairs with a distance of less than 10 kb and frequency of occurrence no less than 10 are extracted from the data to construct the network map. In addition, we also created a map from resistance genes and insertion sequence pairs with a distance of less than 10 kb and the frequency of occurrence no less than 10.

## Results and Discussion

BacAnt is a browser-based platform to annotate DNA sequences, and to visualize the annotation results. When using the web interface, the user first has the choice to upload DNA sequences as Fasta, Seq or GenBank files (Figure 1). The user then has the option to select one or multiple databases, which include ResDB, IntegronDB or TransponsonDB for sequence annotation. After the DNA sequence is submitted, the python-based program BacAnt will identify ARGs and MGEs in the bacterial nucleotide sequence.

**Figure 1.**
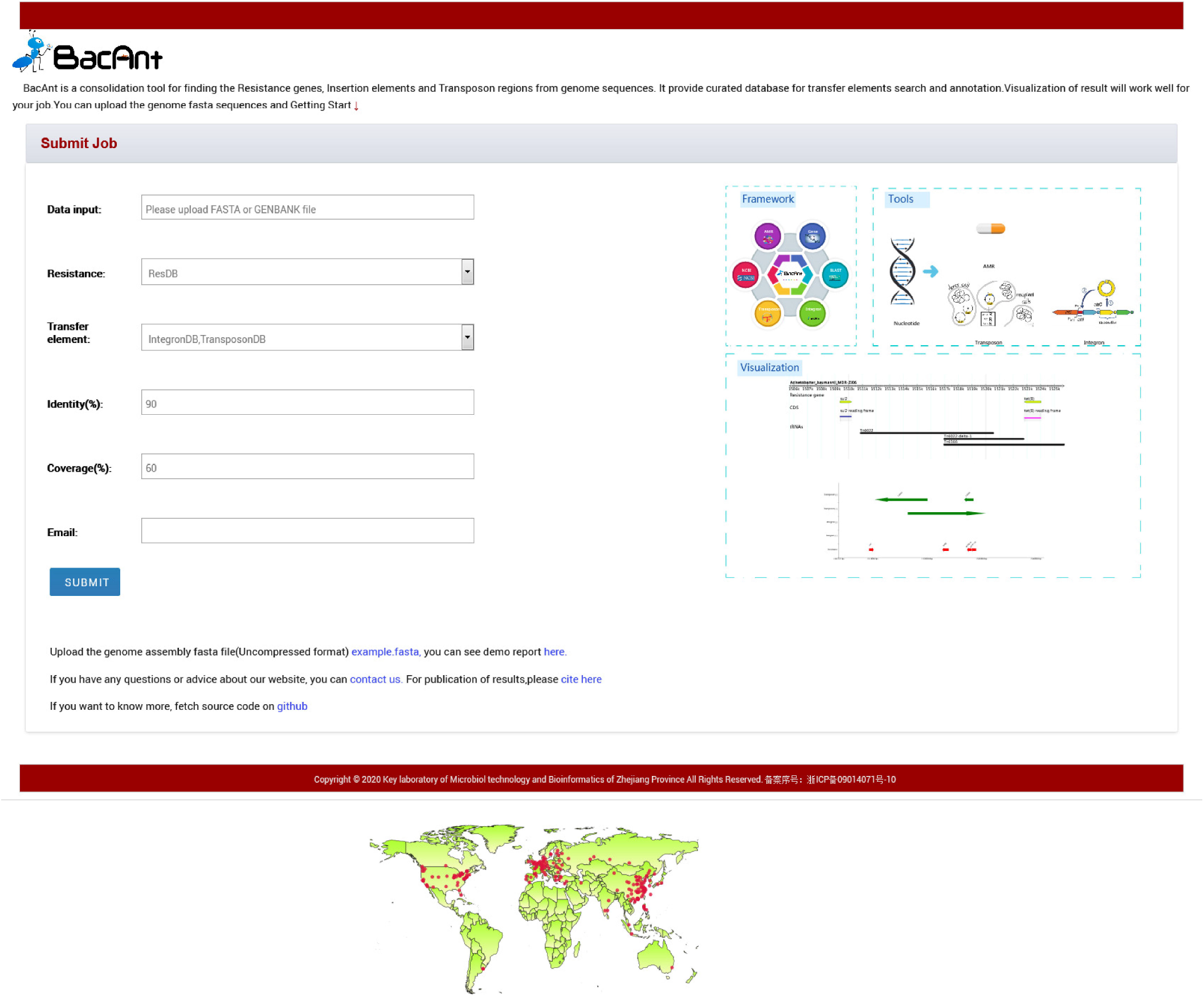
Screenshots of the BacAnt web interface. Users upload an assembled file from their local personal computer and select the desired annotation database. “Framework” lists the public database and tools integrated in BacAnt; “Tools” functions for whole genome sequence annotation based on user uploaded sequence(s); “Visualization” allows to visualize the annotation results for uploaded sequence(s).

The output of BacAnt commences with a summary of the annotation, followed by up to four tables including the annotation result from each database; should they have been selected in the first step. The final part of the BacAnt output visualizes an annotation result which is combined from all three databases. All annotation results obtained by running BacAnt, including figures and genbank files, can then be downloaded. The genebank files generated by BacAnt are compatible with Easyfig (Sullivan et al., 2011). As an example we used *A. baumannii* MDR-ZJ06 (NC_017171.2) to display the result of the annotation by BacAnt (Figure 1). The annotation output of the MDR-ZJ06 strain shows that the isolate harbors 19 resistance genes, 124 integrons and 17 transposons.

BacAnt was validated with NCBI AMRFinder using 1100 selected genomes. The file output “AMR.possible.tsv” in the BacAnt result was used for analysis in NCBI AMRFinder to test which of the programs is able to identify a larger number of ARGs with high accuracy. Both programs reported similar results regarding the number of resistance genes (Table S4). However, the number of ARGS in BacAnt is slightly larger than that of NCBI AMRFinder. Some resistance genes were absent in the output of NCBI AMRFinder: aac(6’)-Iaa(NC_003197,aminoglycoside N-acetyltransferase) in *S. enterica, bla*_EC-15_(NG_049081,class C extended-spectrum beta-lactamase EC-15) in *E. coli*, BcII(NG_056058,BcII family subclass B1 metallo-beta-lactamase in *B. cereus*.

To investigate whether a relationship between ARGs and ISs exists, we extracted the number of ARGs and ISs from the results of the BacAnt analysis of four species genome sequences from NCBI. The results show that the number of ARGs does not correlate with the number of ISs in the four species (Pearson’s R2 < 0.8, Figure 2). Previously, it was reported that at least eight MGEs were detected together with ARGs in *A. baumannii* (Leal et al., 2020). The plasmids were grouped into three categories based on the DNA transfer machinery: conjugative, mobilizable and nonmobilizable (Smillie et al., 2010). Che *et al*. showed that most ARG genes are located in conjugative plasmids, which -together with Insertion Sequences-play the most important role in mediating the horizontal transfer of ARGs (Che et al., 2021). When the relationship between ARG and ISs were investigated, we did not analyze the location of ARGs which might explain why no significant correlation between the two was observed in our study.

**Figure 2.**
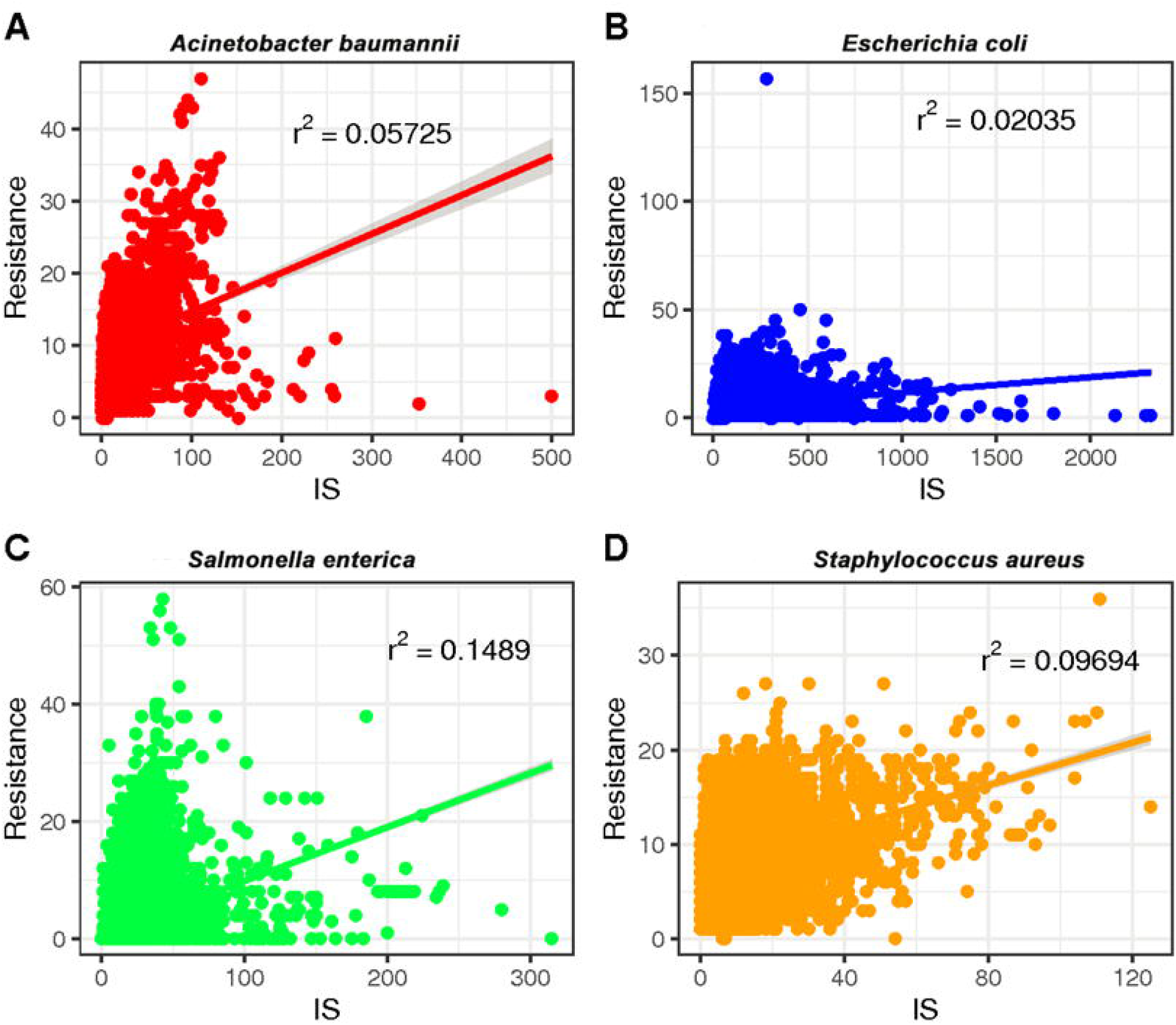
The relationship between the number of drug-resistance genes (ARGs) and insertion sequences (IS) from four species (A: *A. baumannii*, B: *E. coli*, C: *S. enterica*, D: *S. aureus*). The number of ARGs and ISs of each sample were extracted from the results of the BacAnt analysis, and the number was plotted as the horizontal and vertical coordinates respectively. The scatter plot was created using the ggplot2 package in R (V3.6.2), and a trend line generated.

We also calculated the pairwise association between ISs and ARGs. The results were subjected to a permutation test to differentiate between statistically significant associations and random chance (Razavi et al., 2020). Only statistically significant associations (*P* < 0.001) of ISs and ARGs were analyzed (Table S5). We identified commonly found ARG pairs that were in close proximity to the insertion sequence (< 10 kb apart), which allows the detection of gene cassettes that may play an important role in evolution, regulation and ARG exchange. ARG cassettes are generally small (2-7) and specific to the species we investigated (Figure 3). The ARG cassettes for *A. baumannii* and *E. coli* were larger and more stable than the cassettes in *S. aureus*, which is consistent with a previously published observation (Chng et al., 2020). In the case of *A. baumannii*, we identified two ARG cassettes, with one containing the genes *mph(E), msr(E), armA, aadA1, sul1, aac(6’)-Ib* and *catB8*. The ARG cassette which contained *mph(E), msr(E)* and *armA* was described previously (Chng et al., 2020). For *E. coli*, the program identified a stable small cassette that shows overlap with that of *A. baumannii*, including *sul2, aph(3”)-Ib* and *aph(6)-Id*. When genomes of *S. aureus* were analyzed, the program BacAnt found two ARG cassettes; the first one contained the genes *bla*PC1 and *bla*R1, while the second one encoded for *mecR1* and erm(A).

**Figure 3.**
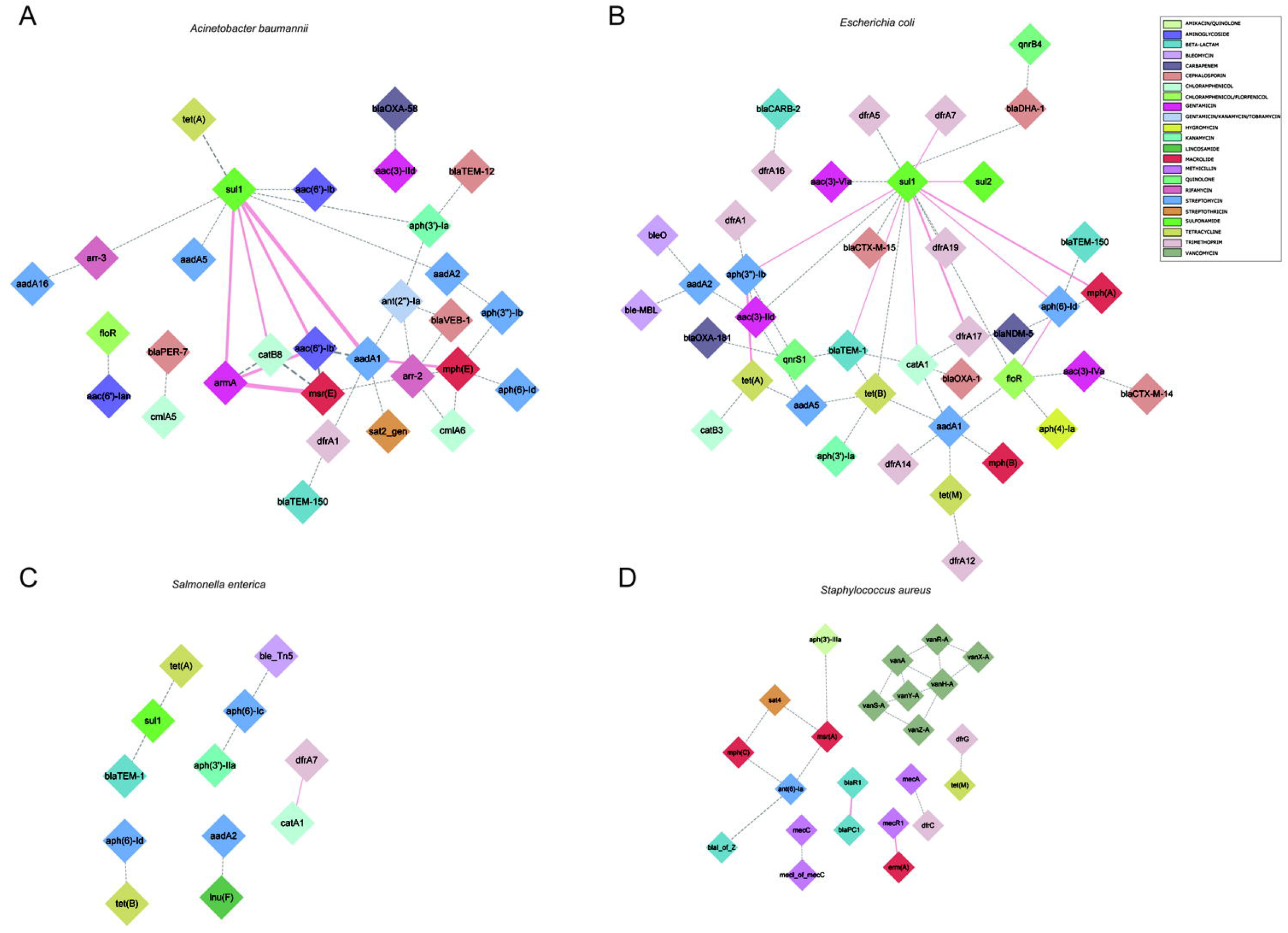
Genomic proximity of antibiotic resistance genes (ARGs) in four species (A: *A. baumannii*, B: *E. coli*, C: *S. enterica*, D: *S. aureus*). ARG pairs with a distance of less than 10 kb and frequency of occurrence no less than 10 were screened to construct the network. Diamonds are used to represent ARGs, solid lines are used to represent gene pairs whose frequency is not less than 80% of the total number and labeled as pink, dotted lines represent gene pairs with frequency less than 80% of the total number and are marked as gray, and the width of the line indicates the frequency. Different antibiotics are displayed in different colors, corresponding to the ARGs.

We also identified commonly found ARG-IS pairs that were in close proximity to the insertion sequence (< 10 kb apart). For the ARG-IS pairs, IS*Vsa3* containing the genes *aph(6)-Id, aph(3”)-Ib* and *tet(B)* comprised the top three ARG-IS pairs in *A. baumannii* (Figure 4A). For ARGs number, IS*26*, IS*Aba1* and IS*Vsa3* were the top three active ISs (Figure 5A). IS*26* was the most abundant IS in *E. coli* and *S. enterica* (Figure 5B, Figure 5C). In *S. aureus, mecA* with diverse IS (including IS*257-3*, IS*431mec*, IS*257R1*, IS*257-1*, IS*257R2*, IS*431L* and IS*431R*) are the top seven ARG-IS pairs (Figure 4D, Figure 5D). The result of this study confirmed that IS*6* family elements IS*26* and IS*257* play an important role in the dissemination of ARGs in *A. baumannii, E. coli, S. enterica* and *S. aureus* (Partridge et al., 2018). As previously reported, we also observed notable differences between important MGEs in *A. baumannii, E. coli, S. enterica* and *S. aureus* (Partridge et al., 2018).

**Figure 4.**
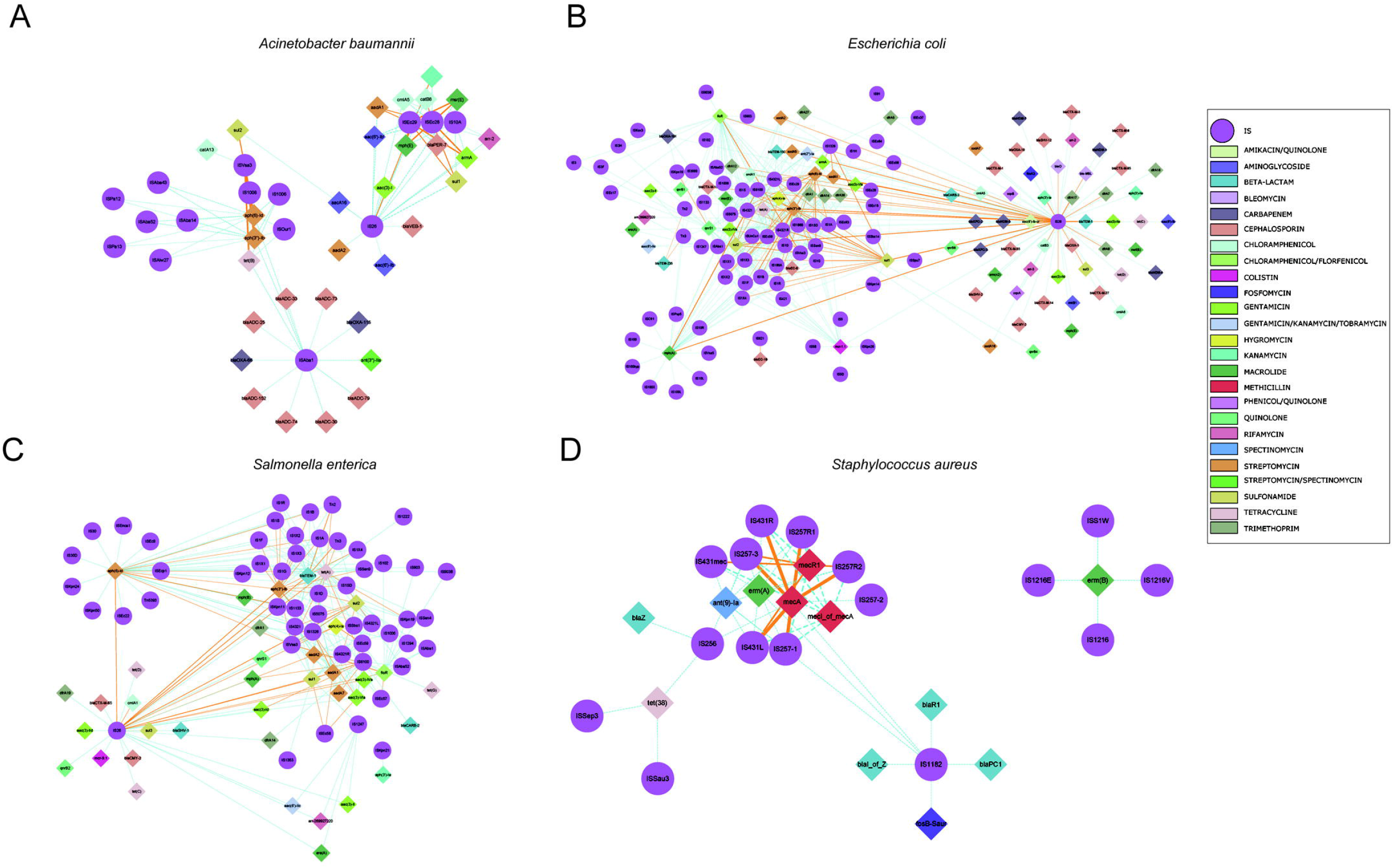
Genomic proximity of antibiotic resistance genes (ARGs) and Insertion sequences (ISs) in four species: A) *A. baumannii*, B) *E. coli*, C) *S. enterica* and D) *S. aureus*. ARGs and ISs pairs with a distance of less than 10 kb and frequency of occurrence no less than 10 were screened to construct the network. Diamond symbols are used to represent ARGs, circles for ISs. The solid line is the gene pair whose frequency is not less than 80% of the total number and labeled as orange, the dotted line represents the gene pair whose frequency is less than 80% of the total number and labeled as sky blue, and the width of the line indicates the frequency. Different colors are assigned to the antibiotics corresponding to the drug-resistant genes, the insertion sequences are displayed in gray.

**Figure 5.**
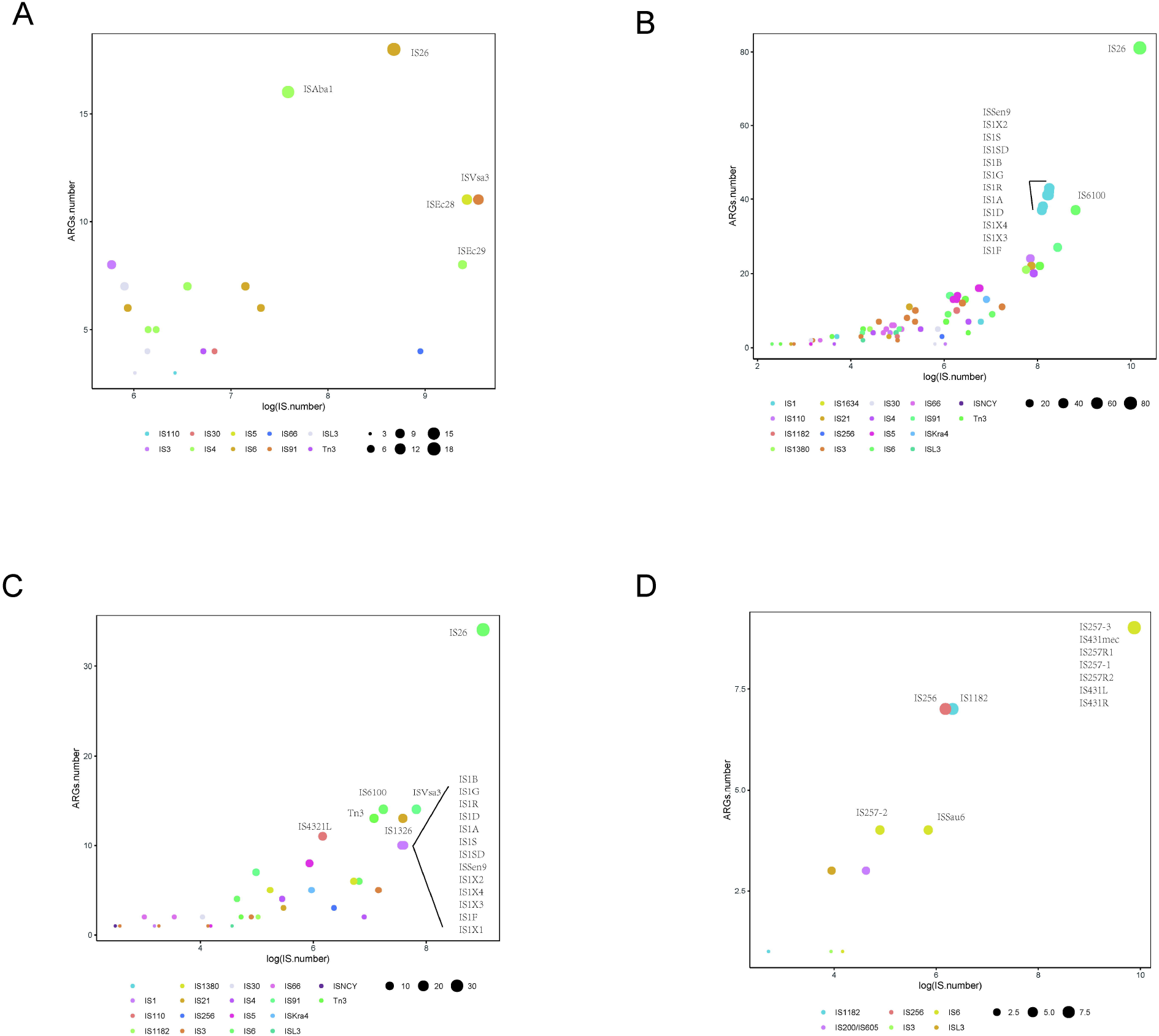
Distribution of Insertion sequences (ISs) with statistically significant association with different types of antibiotic resistance genes (ARGs) within a 10 kb distance. Different colors show various IS, and the size of the circles indicate the presence of their associations with ARGs in A) *A. baumannii*, B) *E. coli*, C) *S. enterica* and D) *S. aureus*.

Using BacAnt, we also explored the relationship between ARGs and transposons. Tn*6292* was the most commonly observed transposon containing ARGs in *A. baumannii, E. coli* and *S. enterica* (Figure 6A, Figure 6B, Figure 6C). In *S. aureus* the most prevalent transposon with ARGs was identified to be Tn*552* (Figure 6D). Tn*6292* belongs to the Tn*3*-family and harbored an IS*26* at the right end (Chen et al., 2020). Tn*6292* also contained a quinolone resistance region *qnrS1* (Feng et al., 2016). Multidrug-resistance bacteria containing Tn*6292* are commonly observed in China (Li et al., 2018), and possibly accelerate the emergence and spread of multidrug-resistant pathogens. Tn*552* belonged to Tn*7* family, comprised of BlaZ, BlaR1 and BlaI proteins. BlaR1 is the sensor protein for the extracellular β-lactam antibiotics. The overproduction of the beta-lactamase BlaZ were responsible of β-lactam resistance. Tn*552*-like element was thought as the origin of the all β-lactamase genes in staphylococci (Gregory et al., 1997).

**Figure 6.**
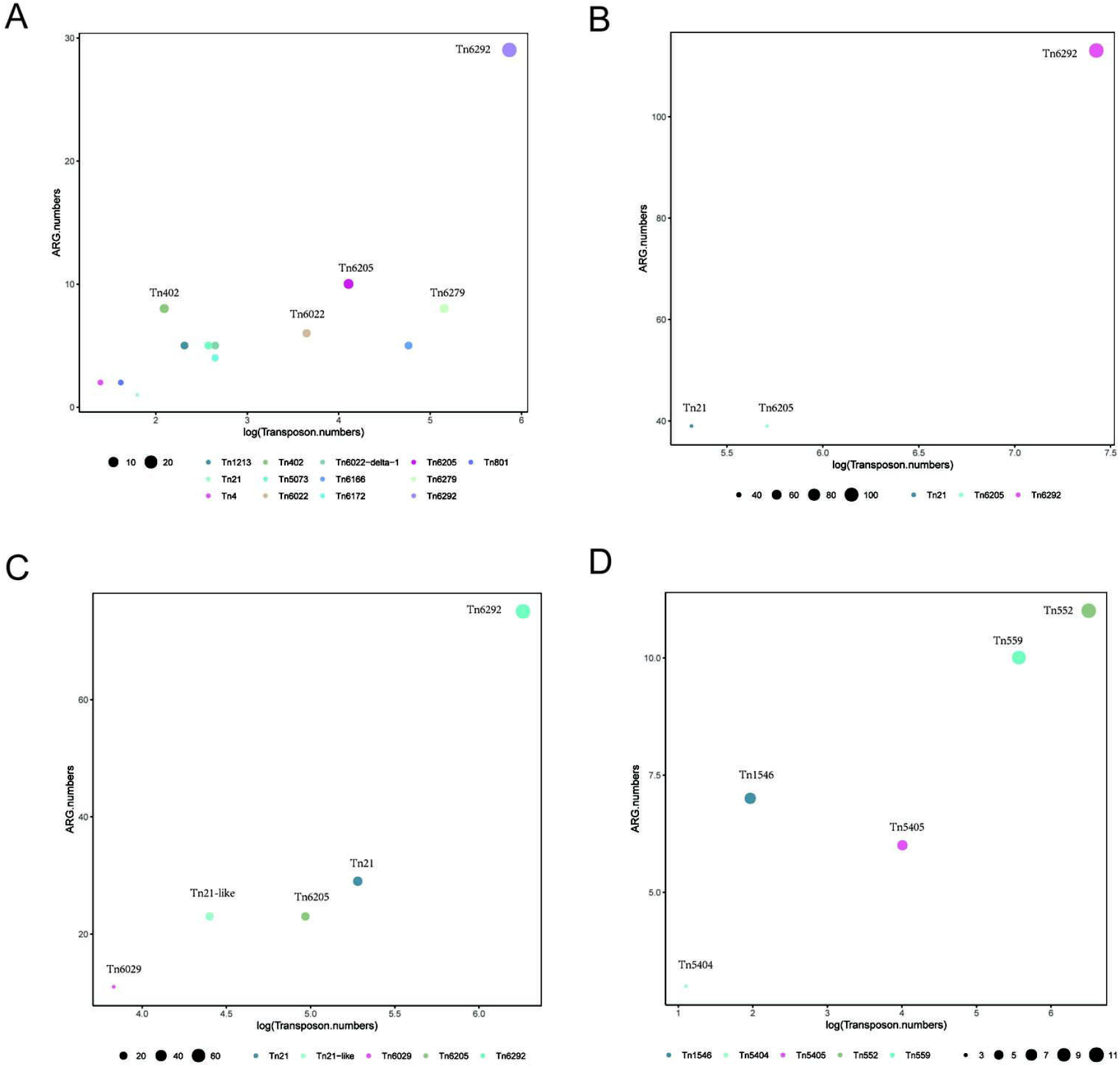
Distribution of transposons with statistically significant association with different types of ARGs within a 10 kb distance. Different colors show various transposons, the size of the circles indicate the presence of their associated ARGs in A) *A. baumannii*, B) *E. coli*, C) *S. enterica* and D) *S. aureus*.

In order to be able to extract the maximum amount of information from whole genome sequence data, we need the improve annotation and analysis methods for MGEs (Partridge et al., 2018). In this work, we created a webserver that is easy to use and allows the annotation of ARGs, ISs, integron, and transposable elements at the same time. The pipeline generates genbank files automatically, which are compatible with easyfig for comparative genomic analysis, which will accelerate the bioinformatics analysis of ARG-related sequences.

## Supporting information

Tabl S2.1

## Data Availability Statement

BacAnt code and standalone software package are available at https://github.com/xthua/bacant with an accompanying web application at http://bacant.net.

## Author Contributions

HC, YY and XH designed the study. QL, MW and WH established the BacAnt. XH, QL, MD, WH, JW and BL analyzed the bioinformatics data. XH, JH, WH and SL wrote the manuscript.

